# Voltage-gated Ca^2+^ influx in murine white fat adipocytes: stimulation by growth hormone but not membrane depolarization

**DOI:** 10.1101/2020.10.27.357699

**Authors:** Nneoma E Akaniro-Ejim, Paul A. Smith

## Abstract

In white fat adipocytes voltage-gated Ca^2+^ channels are constitutively active. Since the adipocyte membrane potential (Vm) is controlled by Cl^−^ we investigated if changes in [Cl^−^]_o_ can affect the activity of voltage-gated Ca^2+^ channels and intracellular calcium, [Ca^2+^]_i_.

Adipocytes were isolated from epididymal fat of CD-1 mice. [Ca^2+^]_i_ was imaged with epifluorescent microscopy at 28°C. Constitutive voltage-gated channel activity was confirmed by the ability of verapamil to decrease [Ca^2+^]_i_.

Substitution of [Cl^−^]_o_ to 113, 53 and 18 mM with the membrane impermeant gluconate anion decreased [Ca^2+^]_i_ from 114±8.7 to 106±7.5, 101±7.1 and 97±6 nM respectively. Substitution of [Cl^−^]_o_ with glutamate mimicked the ability of gluconate to decrease [Ca^2+^]_i_.

To explore if anions affected [Ca^2+^]_i_ via chelation of external Ca^2+^, [Ca^2+^]_o_ was analyzed by potentiometry. Gluconate, glutamate, aspartate and methylsulphonate had Ca^2+^ association constants of 17±1.8, 12±1, 8.2±4.1 and 3.3±0.5 L^−1^ M respectively.

Substitution of 134 mM [Cl^−^]_o_ with gluconate decreased [Ca^2+^]_o_ from 2.6 mM to 200 μM; the effect of this anion on [Ca^2+^]_i_ was mimicked by a decrease of [Ca^2+^]_o_ to 200 μM in standard [Cl^−^]_o_ solution. Conversely, titration of [Ca^2+^]_o_ from 200 μM back to 2.6 mM in 134 mM gluconate solutions abolished the effect of this anion on [Ca^2+^]_i_. Substitution of [Cl^−^]_o_ with methylsulphonate to affect Vm did not affect [Ca^2+^]_i_. Whereas, growth hormone at 10-20 nM increased [Ca^2+^]_i_, an effect blocked by verapamil or absence of [Ca^2+^]_o_. In conclusion, growth hormone, but not changes in Vm, can increase voltage-gated Ca^2+^ channel activity and [Ca^2+^]_i_ in white fat adipocytes.

**Key points:**

- [Ca^2+^]_i_ plays a key role in the metabolic and endocrine functions of white fat adipocytes.
- In adipocytes basal [Ca^2+^]_i_ is maintained by voltage-gated Ca^2+^ channels constitutively active at their resting membrane potential, Vm, which is predominantly controlled by Cl^−^ permeability.
- Substitution of [Cl^−^]_o_ to depolarize Vm with gluconate or glutamate, did not increase but decreased [Ca^2+^]_i_. an action due to chelation of extracellular Ca^2+^. This effect was not seen with methylsulphonate, which did no chelate Ca^2+^ but did not affect [Ca^2+^]_i_.
- Growth hormone elevated, [Ca^2+^]_i_ an effect blocked by inhibitors of voltage-gated Ca^2+^ channels
- In adipocytes, voltage-gated Ca^2+^ channel activity appear recalcitrant to changes in Vm, but are however gated by growth hormone.

## Introduction

Extracellular Ca^2+^-influx is critical to white fat adipocyte (WFA) function. In particular, Ca^2+^-influx is implicated in the processes of fat storage (Arruda & Hotamisligil, 2015): lipolysis (Allen & Beck, 2000) and lipogenesis (Avasthy *et al.*, 1988). Furthermore, the secretion of adipokines, such as adiponectin and leptin, require Ca^2+^-influx for a permissive role in their secretory process (Cammisotto & Bukowiecki, 2004; Komai *et al.*, 2014). Animal and clinical studies suggest that the Ca^2+^ metabolism of white fat adipocytes is associated with the metabolic syndrome (Hvarfner *et al.*, 1988; Ni *et al.*, 1995; Baumbach *et al.*, 2014). Pharmaceutical evidence suggests that Ca^2+^-influx into WFA can occur via L-type voltage-gated calcium channels, VGCCs (Izawa *et al.*, 1983; Draznin *et al.*, 1987; Avasthy *et al.*, 1988; Fedorenko *et al.*, 2019*b*). Moreover, ^45^Ca^2+^ uptake (Martin *et al.*, 1975) and Ca^2+^ imaging studies (Fedorenko *et al.*, 2019*b*) demonstrate that in WFA, Ca^2+^-influx can occur via constitutively active L-type VGCCs. Such observations are consistent with the fact that murine white fat adipocytes have a membrane potential, Vm, of −30mV (Bentley *et al.*, 2014) a value within the activation gating range of L-type VGCCs (Koschak *et al.*, 2001). The depolarised Vm of WFA originates from a predominant Cl^−^ permeability of the adipocyte plasma membrane; consequently the Vm of WFA behaves as a Cl^−^ electrode in response to changes in extracellular Cl^−^ concentration, [Cl^−^]_o_(Bentley *et al.*, 2014). Given these facts, the aim of this study was to investigate if the [Ca^2+^]_i_ of WFA could be affected by alterations in VGCC activity mediated by changes in [Cl^−^]_o_ and Vm.

## Materials and Methods

### Ethical approval

All animal care and experimental procedures were carried out in accordance with the UK Home Office Animals (Scientific Procedures) Act (1986). Animals were killed by Schedule 1 cervical dislocation followed by decapitation. All animal procedures were approved by the local ethical committee (ASPA 000187, BMU, School of life Sciences, Nottingham).

### Isolation and preparation of adipocytes

White fat adipocytes were isolated from the epididymal fat pads of male CD-1 mice (fed *ad libitum,* 12 hr. dark/light cycle, weight 25 – 35 g; Charles River Laboratory, Kent, UK). Adipocytes were isolated by collagenase digestion as previously described (Bentley *et al.*, 2014).

### Solutions

Unless stated otherwise, adipocyte isolation and experiments were performed in Hank’s buffer solution composed of (in mM): NaCl 138, NaHCO_3_ 4.2, KCl 5.6, MgCl_2_ 1.2, CaCl_2_ 2.6, NaH_2_PO_4_ 1.2, HEPES 10 (pH 7.4 with NaOH, calculated ionic strength 166 mM), glucose 5 and 0.01% wt./vol BSA. For Ca^2+^ free solutions, CaCl_2_ was replaced with equimolar MgCl_2_. Unless stated otherwise drugs were from Sigma, Poole, UK and chemicals were of analytical grade. Verapamil was used at a relatively high concentration of 20 μM for reasons discussed elsewhere (Fedorenko *et al.*, 2019*b*).

### Measurement of intracellular free calcium concentration, [Ca^2+^]_i_

Intracellular free calcium concentration of the adipocytes, [Ca^2+^]_i_ was measured with methods similar to those previously described for rat (Fedorenko et al 2019). Briefly, primary adipocytes were attached to poly-L-lysine (25-100 μg ml-1) coated coverslips and incubated with the Ca^2+^ fluorophore Fluo-4 AM (1 μM; Molecular Probes) in modified Hank’s solution for 1 hour at 21-23°C in the dark. Coverslips were mounted in a perifusion chamber on an Axiovert 135 Inverted microscope equipped for epifluorescence (Carl Zeiss Ltd, UK). Cells were focused to maximize equatorial circumference and epifluorescence. Adipocytes were identified under Kohler illumination as 50-100 μm diameter spheroids with a nuclear protuberance. Experiments were performed 1-4 hour post isolation.

Fluo-4 was excited at 450-490 nm, the emitted light band pass filtered at 515-565 nm. Images were captured at 1 Hz with a Coolsnap HQ2 camera (Photometrics, Tucson, AZ, USA) with Imaging Workbench (Ver. 6.2, INDEC Biosystems: RRID:SCR_016589). Cells were continuously perifused at 28°C in Hanks solution. In basal, control solutions, [Cl^−^]_o_ was 152 mM. The effect of extracellular Cl^−^ removal on [Ca^2+^]_i_ was explored by equimolar substitution of bath NaCl with sodium salts of organic anions.

To measure [Ca^2+^]_i_, a region of interest (ROI) was drawn around each cell, background corrected and the time course of its overall mean fluorescent intensity measured. Fluorescence was then calibrated by a two-point method (Ni *et al.*, 1994): the maximum fluorescence value, Fmax; determined by permeabilization of the cells with Triton X-100 (0.0125-0.1%) followed by washout of the dye to determine the minimum, Fmin. [Ca^2+^]_i_ was calculated with the following equation:

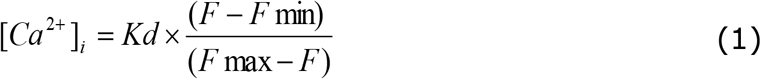

Where F is the background corrected fluorescence, and Kd the dissociation constant of Fluo-4: 345 nM (Bentley *et al.*, 2014). Data was only used from cells in which [Ca^2+^]_i_ was calibrated and stable; the latter established as a fluorescent signal that did not continually increase or decrease, regardless of intervention, during the time course of experiment.

To visualize spatial variation of [Ca^2+^]_i_ in the few cells that responded to growth hormone it was necessary to mitigate heterogeneity in fluorescence that arose through uneven dye loading and differences in cytoplasmic volume. Since these parameters affect Fluo-4 fluorescence in a proportional manner, the ratio of the fluorescence signal in the presence of treatment with that in its absence was undertaken using ImageJ2 (RRID:SCR_003070) with floating point arithmetic (Rueden *et al.*, 2017).

### Measurement of free Ca^2+^ in the extracellular solution

The free calcium concentration of solutions, [Ca^2+^]_o_, were measured with a combination Ca^2+^ electrode (EDT direct ion, Dover, UK). The electrode was calibrated with three points: 25, 2.5 and 0.25 mM CaCl_2_ at 20-22°C. Calibration solutions were made by serial dilution of a 1 M CaCl_2_ standard with a 150 mM NaCl, 10 mM HEPES (pH 7.4 with NaOH) solution; ionic strengths, I, were calculated as 230, 163 and 156 for 25, 2.5 and 0.25 mM CaCl_2_ respectively. Resultant electrode voltages, E_Ca_, were fit with the Nernst equation for Ca^2+^ and the [Ca^2+^] for test solutions obtained by extrapolation. To check for electrode voltage drift, calibration was performed both prior to and post measurement, although in practice drift was immeasurable. The electrode was insensitive to Mg^2+^ ions as E_Ca_ was unaffected by addition of 4.8 mM MgCl_2_ to the Ca^2+^ calibration solutions

### Estimation of anion affinity for Ca^2+^

At the physiological pH of 7.4, all four anions tested: gluconate, glutamate, aspartate and methylsulphonate primarily exist as monovalent anions with the zwitterionic amino and carboxylic acid dipoles undergoing mutual charge neutralization (Chemicalize.com). In the presence of Ca^2+^, the anion ligand can exist in three forms: the free ligand, L^−^; as the monomolecular complex, CaL^+^; and as the bimolecular complex, CaL_2_. These are shown in the following reaction scheme, where K_a1_ and K_a2_ represent the appropriate association constants (L M^−1^) for the monomolecular and bimolecular complex respectively:

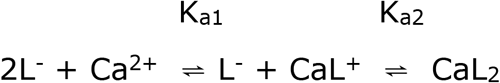

Where at steady state

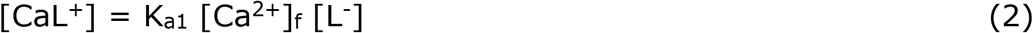

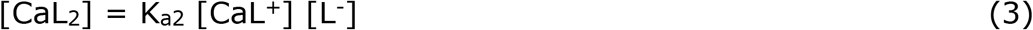

and

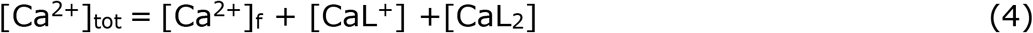

Substitution of equations 2 and 3 into 4 followed by rearrangement gives

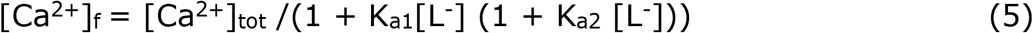

Where [Ca^2+^]_tot_ is the total concentration of Ca^2+^ and [Ca^2+^]_f_ the concentration of free Ca^2+^; [L^−^] is assumed in excess such that its concentration is unaffected by chelation. Three possible variants of this model exist: where binding of the second ligand is influenced by bound ligand such that K_a1_ > 0 and K_a2_ > 0 and K_a1_ ≠ K_a2_; where binding of the second ligand is similar to, but independent of, bound ligand K_a1_ > 0 and K_a1_ ≈ K_a2_,; where binding of the second ligand is prevented by bound ligand K_a1_ > 0 and K_a2_ → 0. The equation for the latter case simply being:

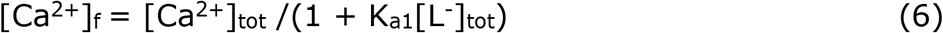

In the case where [L^−^] > [Ca^2+^]_tot_, [CaL^+^] can be estimated by:

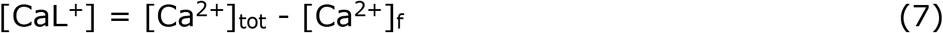

Or to account for changes in [L^−^] due to Ca^2+^ chelation:

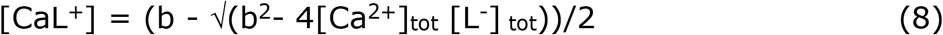

Where K_a2_ → 0 and b = [Ca^2+^]_tot_ + [L^−^]_tot_ +1/K_a1_ with

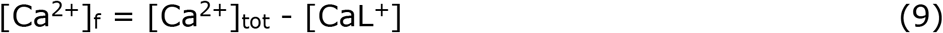

and

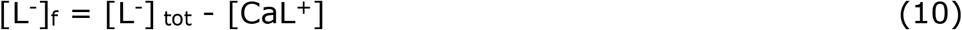

The models described by equations 5 and 6 were fitted for each set of data with the best fit model determined by the F-test.

At the low ionic strengths, I, employed in this study ideality of behaviour was assumed and ion concentration was used instead of activity, γ.

### Experimental Determination of K_a1_ and K_a2_

Hank’s solutions in which 134 mM NaCl were substituted with a Na^+^ salt of the test anion were prepared. These solutions were mixed ratiometrically with 152 mM Cl^−^ Hank’s solution to form solutions of known anion composition of [L^−^] from 0 to 134 mM; that is a [Cl^−^] from 152 to 18 mM. For each of these solutions, [Ca^2+^]_o_ was measured and plotted as a function of [L^−^] (Fig. 2). To determine K_a1_ and K_a2_ this relationship was fitted with equation 5 by a least squares algorithm here [Ca^2+^]_tot_ was constrained to the value of [Ca^2+^]_o_ measured in the absence of L-; median value 2.47 mM (2.36-2.5; 95% C.I., n = 17). Since solutions also contain Mg^2+^, a decrease in free ligand concentration [L^−^] is expected via Mg^2+^ binding. To ascertain if Mg^2+^ binds to [L^−^] and affects [Ca^2+^]_o_ experiments were repeated with solutions in which MgCl_2_ was omitted; this did not significantly affect the values of K_a1_ and K_a2_. Secondly, if it is assumed that the same reaction scheme occurs for Mg^2+^ as for Ca^2+^, but the association constants for Mg^2+^≫ Ca^2+^, which maybe expected based on anion binding to the alkaline earth elements (Lumb & Martell, 1970), then a decrease in [L^−^] of 2.4 mM at most is expected. To test this idea, data for solutions in which Mg^2+^ was already present were fit with equation 5 but with a value of [L^−^] decremented by 2.4 mM; this procedure similarly had little impact on K_a1_ and K_a2_. We did not attempt to derive the on K_a1_ and K_a2_ for Mg^2+^ binding to the ligands, nor did we explore the effects of lowered extracellular Mg^2+^ on [Ca^2+^]_i_ in adipocytes. In order to directly test the effect of [Ca^2+^]_o_ on [Ca^2+^]_i_ when the anion solutions chelated Ca^2+^, Hank’s solutions were used in which the [Ca^2+^]_o_ was titrated back to the required concentrations as measured with a Ca^2+^-electrode.

### Simulation of chelation

To check our Ka values for gluconate, the association of Ca^2+^ with ligand was simulated via a multicompartment model (Modelmaker Ver 4.0, AP Benson). K_a_ was taken as the value determined by Ca^2+^ measurements described (Table 1) and the effect of incremental substitution on free Ca^2+^, [Ca^2+^]_f_ was simulated. The resultant steady-state values for [Ca^2+^]_f_ were then plotted against the different [L^−^]_tot_ as well as [L^−^]_f_, as calculated by the simulation, and the Ka values reextracted by the procedures detailed above.

**Table 1.** Association constants for the Ca^2+^ ligand salts. Data presented as means ± S.D.

| **Anion** | $K_{a1}$ ($\text{L M}^{-1}$)<br>Using $[\text{L}^-]_{\text{tot}}$ | $K_{a2}$ ($\text{L M}^{-1}$) | $K_a$ ($\text{L M}^{-1}$)<br>Using $[\text{L}^-]_f$ | n |
| --- | --- | --- | --- | --- |
| Gluconate | $17.2 \pm 1.8$ | $= K_{a1}$ | | 4 |
| Aspartate | $11.8 \pm 1$ | 0 | $12 \pm 1$ | 3 |
| Glutamate | $8.2 \pm 4.1$ | 0 | $8.4 \pm 4.2$ | 5 |
| Methylsulphonate | $3.34 \pm 0.5$ | 0 | $3.36 \pm 0.5$ | 5 |

### Estimation of Vm

The membrane potential of adipocytes expected on substitution of bath Cl^−^ werecalculated using the Goldman Hodgkin Katz (GHK) constant field equation and parameters as previously published for murine epididymal adipocytes (Bentley *et al.*, 2014).

### Statistical analysis

Data distributions were checked for normality with the D’Agostino & Pearson omnibus test, appropriate inferential tests are given in the text. When the number of different adipocytes, n, was greater than 30 data is shown graphically as box and whisker plots with 10-90-% confidence intervals. When n <= 30 data are shown as scatter plots overlaid with the median and 25%-75% interquartile range. Numerical data are quoted either as the mean ± S.D. or median with 5 to 95% confidence intervals (95% C.I.) to 3 significant figures, where n is the number of cells, f, the number of individual coverslips pooled from up to 5 different animal preparations. We have already commented upon the observation that variation in [Ca^2+^]_i_ for adipocytes with an animal is sometimes greater than between animals (Fedorenko *et al.*, 2019*b*). For this reason, we performed the majority of experiments as repeated measurements with internal controls where possible. When we looked for differences between data collected from different animals, to mitigate inter-animal variation we used percentage values relative to basal for each determination. To determine statistical significance between groups Friedman’s was used with Dunn’s multiple comparison test. Fits of equations to data used a least squares algorithm with the parameters given in text. Comparison between models was made with a sum-of-squares F test. Statistical analyses were performed using Graphpad PRISM (RRID:SCR_002798). Data were considered statistically significant different when p < 0.05 and in graphics is flagged as *, ** when P <0.01, *** when P <0.001 and **** when P <0.0001.

## Results

### Substitution of bath Cl^−^ with gluconate reversibly decreased [Ca^2+^]_i_

Figure 1A shows that substitution of extracellular Cl^−^ with gluconate with the same protocol previously performed to explore the effect of this anion on adipocyte Vm (Bentley *et al.*, 2014) decreased [Ca^2+^]_i_. For the majority of cells tested only a partial recovery of [Ca^2+^]_i_ was observed on return to normal [Cl^−^]_o_ (Figs. 1B, C). Substitution of [Cl^−^]_o_ with gluconate significantly decreased [Ca^2+^]_i_ at all concentrations tested (Fig. 1B). The relative decrease in [Ca^2+^]_i_ magnitude with 18 mM [Cl^−^]_o_, Δ[Ca^2+^]_i_, was negatively correlated (Spearman r = −0.43, p=0.002, n=48) with basal [Ca^2+^]_i_ (Fig. 1D): cells that possessed an initial higher basal [Ca^2+^]_i_ displayed the largest decrease in [Ca^2+^]_i_ when [Cl^−^]_o_ was dropped from 152 to 18 mM. Cells that possessed the largest initial basal [Ca^2+^]_i_ also recovered the least on reinstatement of normal [Cl^−^]_o_ (Fig. 1C;Spearman r = −0.33, p = 0.02, n=48); for these latter cells, [Ca^2+^]_i_ recovered towards the lower end of the initial basal distribution (Fig. 1E).

**Figure 1.**
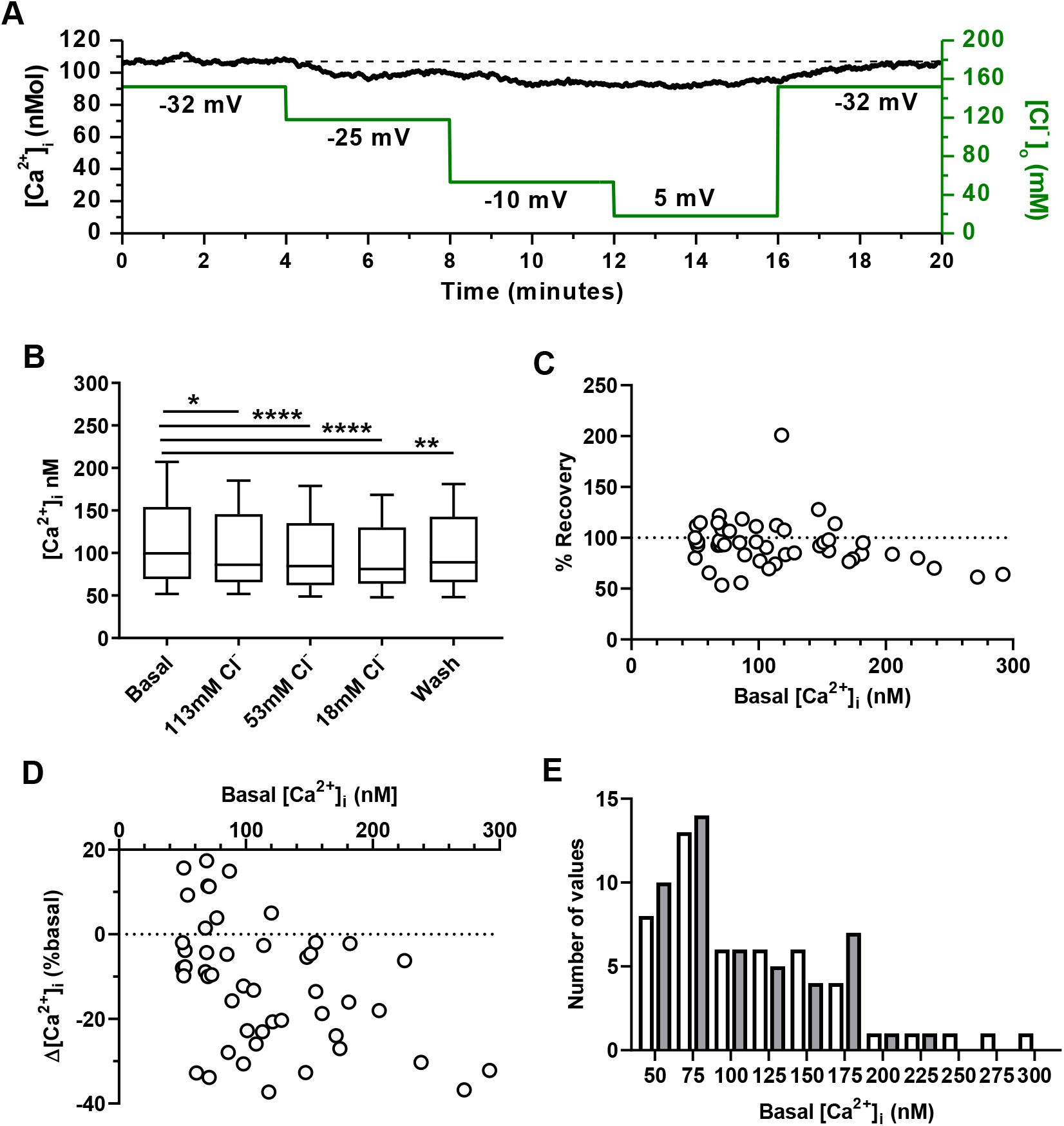
Substitution of bath Cl^−^ with gluconate reversibly decreases intracellular [Ca^2+^] of white fat adipocytes. A) Mean intracellular Ca^2+^ of 5 adipocytes within a single field of view in response to equimolar replacement of bath [Cl^−^]_o_ with gluconate as indicated by the green line. Numbers on graph are Vm values predicted for the various [Cl^−^]_o_ as described in methods. B) [Ca^2+^]_i_ under the conditions indicated as in A) when bath Cl^−^ is substituted with gluconate. In basal and wash Cl^−^ is not substituted and [Cl^−^]_o_ is 152 mM (n = 46, f = 10). Statistical inference by Friedman with Dunn’s multiple comparison tests. C) Relationship between the percentage recovery in [Ca^2+^] on return of [Cl^−^]_o_ from 18 mM to 152 mM and basal [Ca^2+^]_i_. D) Relationship between the decrease in [Ca^2+^]_i_, Δ[Ca^2+^]_i_, observed in 18mM [Cl^−^]_o_ and basal [Ca^2+^]_i_. E) frequency histogram of basal [Ca^2+^]_i_ before (filled bars) and after wash out of gluconate: wash recovery (open bars).

### Organic anions chelate [Ca^2+^]_o_

We previously predicted that removal of extracellular Cl^−^ would depolarize Vm and elicit an increase in voltage-gated Ca^2+^ influx (Bentley *et al.*, 2014); however, this phenomenon was not observed here (Fig. 1.) Since gluconate is reported to buffer Ca^2+^ (Woehler *et al.*, 2014), the possibility exists that the decrease in [Ca^2+^]_i_ we observed when [Cl^−^]_o_ was removed was actually due to a direct drop in [Ca^2+^]_o_ like that previously described when bath Ca^2+^ is substituted with Mg^2+^ (Fedorenko *et al.*, 2019*b*). Although association constants for Ca^2+^-gluconate have been reported, these measurements were either done in low-ionic strength, <0.08M (Skibsted & Kilde, 1972), or under non-physiological conditions (Cannan & Kibrick, 1938; Schubert & Lindenbaum, 1952; Vavrusova *et al.*, 2013). Consequently, we measured the association constant for gluconate in Hank’s solution at physiological ionic strengths 0.17 M and pH 7.4.

Figure 2A demonstrates that substitution of bath Cl^−^ with gluconate monotonically decreased [Ca^2+^]. The relationship between [Ca^2+^] and [Gluconate^−^] was readily described by equation 5 with a 2:1 stoichiometry and similar valued association constants, K_a1_ and K_a2_ (Table 1). On substitution of [Cl^−^] with aspartate, glutamate or methylsulphonate, best fits of [Ca^2+^]_o_ were all achieved with a 1:1 stoichiometry. The overall rank order for association with Ca^2+^ was gluconate > aspartate > glutamate > methylsulphonate; as indicated in Figure. 2 and Table 1. Correction for loss of total ligand, [L^−^]_tot_, by Ca^2+^ chelation gave fractionally larger values of K_a_ with equation 8 (Table 1), with larger differences as K_a_ and the associated chelation increased.

**Figure 2.**
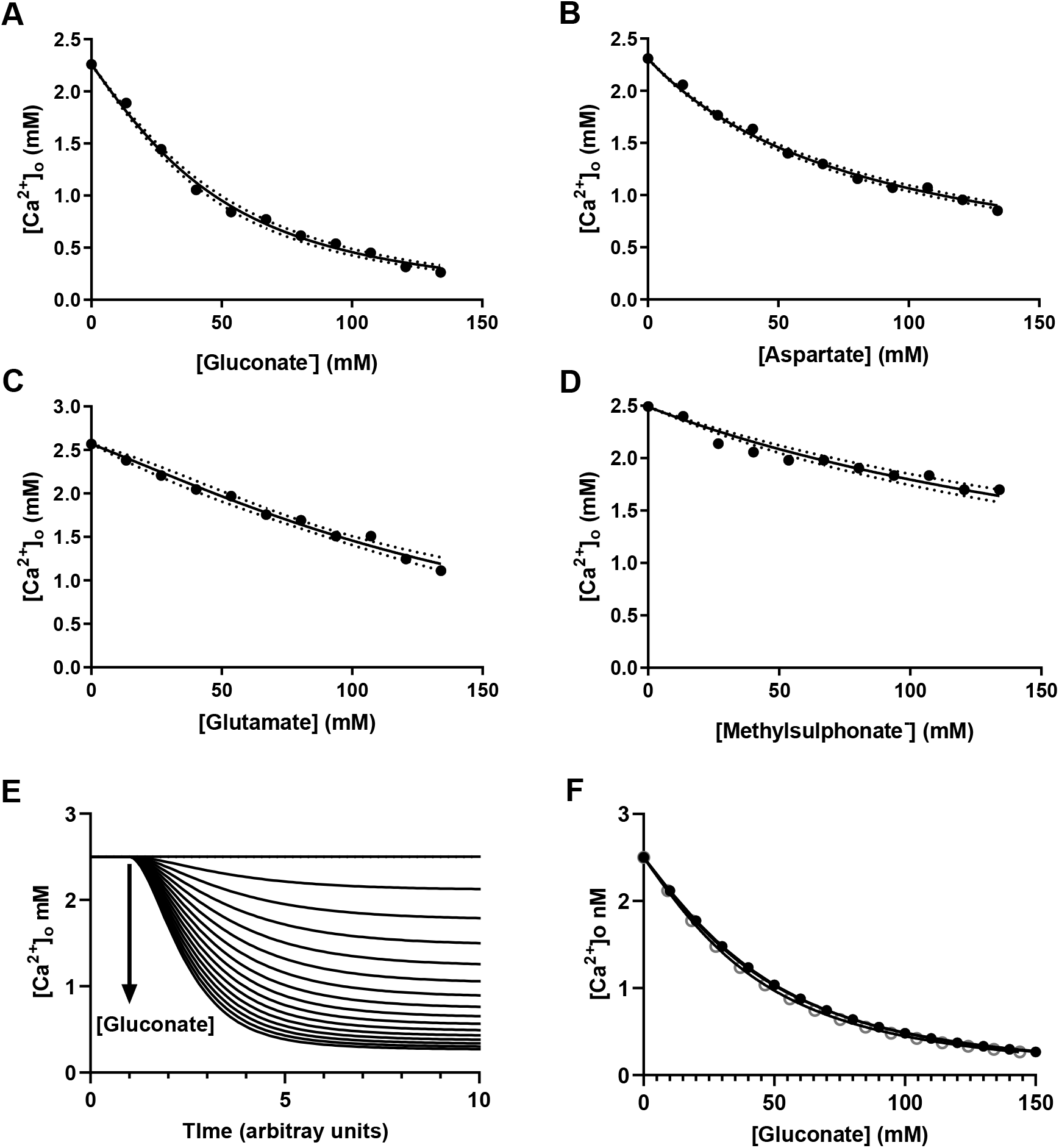
Organic anions chelate extracellular Ca^2+^. A-D) Representative plots of the free [Ca^2+^] against concentration of anion as indicated on equimolar substitution of bath Cl^−^. Each data point is a single determination. Solid lines are fits of equation 5 as indicated in the text. Dotted lines are the 95% confidence intervals for the fit shown with A) K_a1_ = 15.5 L M^−1^ B) K_a1_ = 11.7 L M^−1^, C) K_a1_= 7.3 L M^−1^ D) K_a1_= 3.9 L M^−1^. E) Computed simulation of Ca^2+^ chelation and [Ca^2+^]_f_ in a solution where [Cl^−^] is substituted with gluconate from 0 to 150 mM in 10 mM increments with a 2:1 stoichiometry and a K_a_ of 0.017 L M^−1^ (table 1). Note the decrease [Ca^2+^]_o_ with increase in [gluconate] which is added at arrow. F) Plot of steady state [Ca^2+^]_o_ from the simulation shown in E) against free, ○, and total, •, gluconate concentrations. Solid lines are fits of equation 5 with K_a_ of 0.017 and 0.016 L M^−1^ respectively.

Computed simulation of Ca^2+^ chelation by gluconate (Fig. 2E) yielded identical values of K_a_ when [Ca^2+^]_f_ was fitted for [L^−^]_f_, whereas a slightly lower value of K_a_ was obtained when [L^−^]_tot_. This difference in K_a_ demonstrates that unaccounted loss of free ligand by Ca^2+^ chelation leads to an underestimation of K_a_ as already shown experimentally above.

### Further exploration of the effects of [Ca^2+^]_o_ chelation on [Ca^2+^]_i_

To explore if the decrease in adipocyte [Ca^2+^]_i_ observed with gluconate wasin fact due to chelation of extracellular Ca^2+^ by the organic anion, two approaches were undertaken. First, the effect of a 152 mM [Cl^−^]_o_ Hank’s solution but in which [Ca^2+^]_i_ was decreased to 200 μM, a value similar to that measured in 134 mM [Gluconate^−^] was tested (Fig. 3A). This procedure mimicked the ability of 134 mM [Gluconate-] to lower [Ca^2+^]_i_ (Figs. 3A & B; Friedman, n=15). Secondly, the effect of extracellular Ca^2+^ supplementation by titration to replace that chelated by gluconate was explored. Subsequent substitution of extracellular Cl^−^ with gluconate^−^ but with a solution where [Ca^2+^]_o_ was similar to that measured in normal 152 mM [Cl^−^]_o_ did not affect [Ca^2+^]_i_ (Figs 3C & D; Friedman, n = 16).

**Figure 3.**
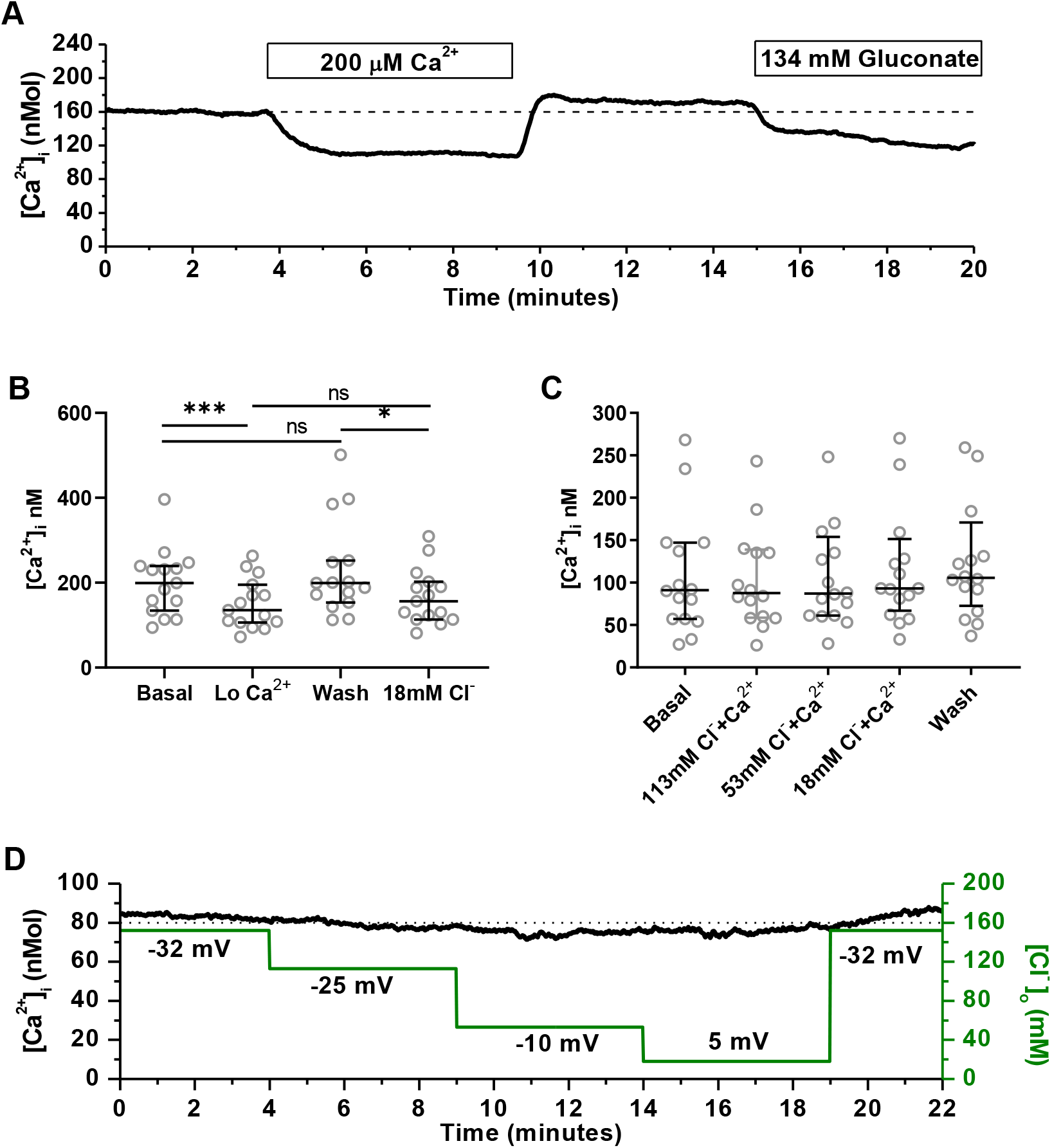
Gluconate decreases intracellular [Ca^2+^] of white fat adipocytes an effect prevented by extracellular Ca^2+^ addition. A) Mean intracellular Ca^2+^ for 4 adipocytes in response to a reduction in bath Ca^2+^, followed by substitution of 134 mM bath Cl^−^ with gluconate. Cells are all within the same field of view. B) [Ca^2+^]_i_ under the conditions indicated as shown in A) (n = 15, f = 4). Statistical inference by Friedman with Dunn’s multiple comparison tests. C) [Ca^2+^]_i_ under the conditions indicated as for D) where bath Cl− is sequentially decreased by substitution with gluconate but bath [Ca^2+^] is maintained at 2.5 mM (n = 16, f = 3). Basal and wash are 152 mM [Cl^−^]_o_. Statistical inference by Friedman with Dunn’s multiple comparison tests. D) Mean intracellular Ca^2+^ for 4 adipocytes in response to equimolar replacement of bath [Cl^−^]_o_ with gluconate, as indicated by the green line, but in which the free [Ca^2+^] is preserved at 2.5 mM. Numbers on graph are predicted Vm values for the various [Cl^−^]_o_ as described in methods. Cells are all within the same field of view.

Since glutamate was also found to chelate [Ca^2+^]_o_ (Table 1), we expected it to act like gluconate and decrease [Ca^2+^]_i_ when substituted for [Cl^−^]_o_. The data shown in Figures 4A & B confirm this notion, where sequential substitution of extracellular [Cl^−^]_o_ by glutamate decreased [Ca^2+^]_i_ in an incremental manner. Both the decrease in [Ca^2+^]_i_, Δ[Ca^2+^]_i_ observed with 134 mM [Glutamate-] 18 mM [Cl^−^]_o_ as well as its subsequent recovery in 152 mM [Cl^−^]_o_ were negatively correlated with the basal level of [Ca^2+^]_i_; Spearman r =−0.52 (p=0.04, n=16; Fig. 4C) and Spearman r = −0.57 (p=0.02, n=16; Fig. 4D) respectively. Since methylsulphonate had the smallest association constant of the four anions tested (Table 1), we expected it to chelate extracellular Ca^2+^ the least and cause the smallest decrease of [Ca^2+^]_i_. This notion is confirmed in Figures 4E & F where substitution of extracellular Cl^−^ with the methylsulphonate anion had no detectable effect on adipocyte [Ca^2+^]_i_ even at 134 mM with 18 mM [Cl^−^]_o_. Figure 4G compares the percentage decreases in basal [Ca^2+^]_i_, Δ[Ca^2+^]_i_, observed with different anions at a concentration of 134 mM with 18 mM [Cl^−^]_o_ compared to those seen in the wash control at equivalent time points. Consistent with our previous findings, only gluconate and glutamate significantly decreased [Ca^2+^]_i_. Although aspartate chelates [Ca^2+^]_o_ with an association constant larger than that for glutamate, this was not tested due to the effects already seen with glutamate being comparable with that of gluconate as well as the previous observation that this anion had partial permeability in Cl^−^ channels for this cell type (Pulbutr *et al.*, 2007).

**Figure 4.**
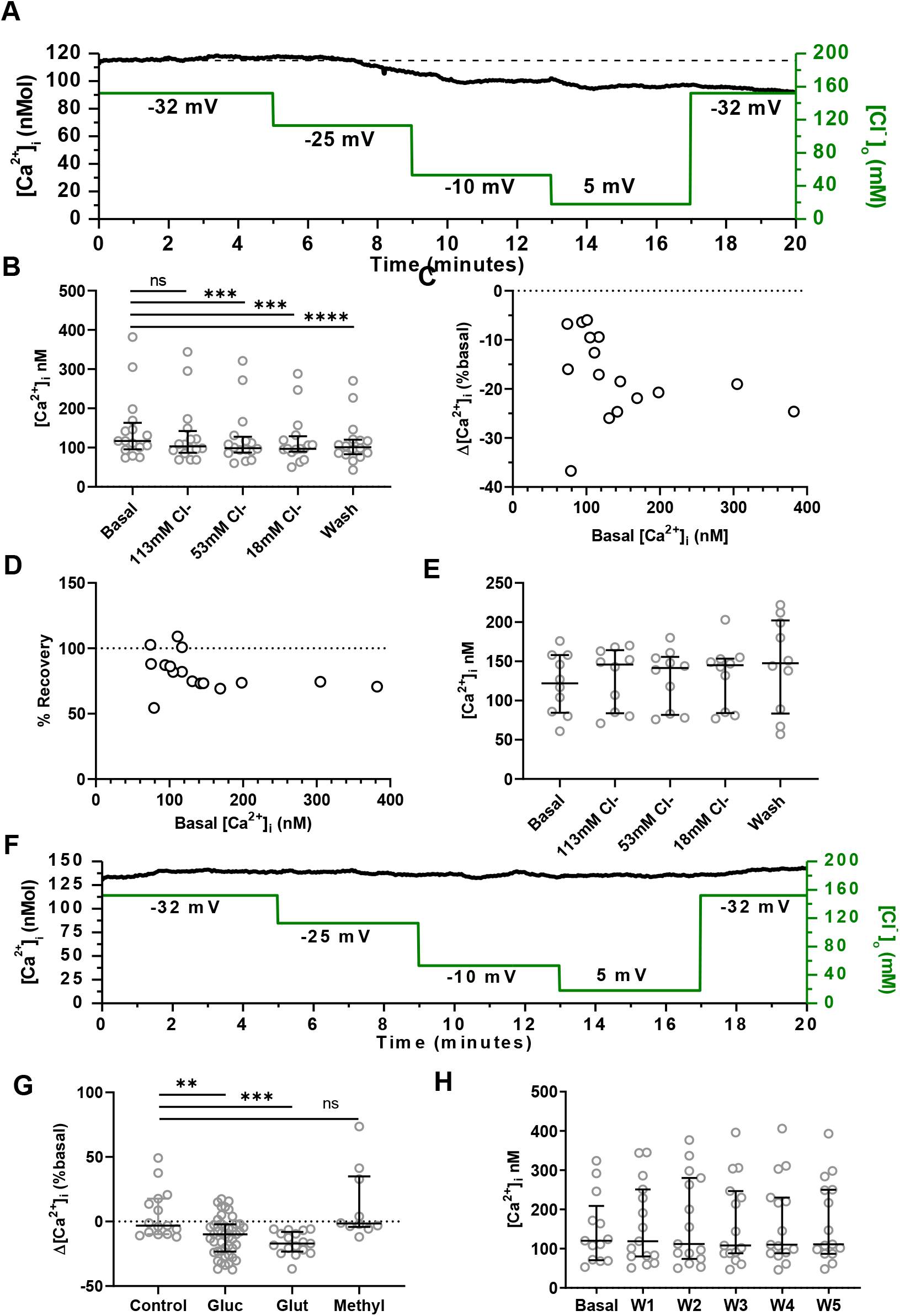
Glutamate, but not methylsulphonate, decreases intracellular [Ca^2+^] of white fat adipocytes. A) Mean intracellular [Ca^2+^] for 5 adipocytes within a single field of view in response to sequential equimolar replacement of bath [Cl^−^]_o_ with glutamate as indicated by the green line. Numbers on graph are predicted Vm values for the various [Cl^−^]_o_ as described in methods. B) [Ca^2+^]_i_ under the conditions shown in A) when Cl^−^ is substituted with glutamate. Basal and wash is 152 mM [Cl^−^]_o_ (n = 16, f = 4). Statistical inference by Friedman with Dunn’s multiple comparison tests. C) Relationship between the decrease in [Ca^2+^]_i_, Δ[Ca^2+^]_i_, observed in 18mM [Cl^−^]_o_ and basal [Ca^2+^]_i_. D) Relationship between the percentage recovery in [Ca^2+^]_i_ on return of [Cl^−^]_o_ from 18 mM to 152 mM and its initial basal value. E) [Ca^2+^]_i_ conditions where Cl^−^ is sequentially substituted with methylsulphonate as shown in F). Basal and wash are 152 mM [Cl^−^]_o_ under the conditions indicated (n = 16, f = 3). Statistical inference by Friedman with Dunn’s multiple comparison tests. F) Mean intracellular Ca^2+^ for 4 adipocytes in response to equimolar replacement of bath [Cl^−^]_o_, as indicated by the green line, with methylsulphonate Numbers on graph are predicted Vm values for the various [Cl^−^]_o_ as described in methods. Cells are all within the same field of view. G) [Ca^2+^]_i_ when 134 mM bath Cl^−^ is substituted with anion shown. Control, no substitution (n = 15); Gluc, gluconate (n = 46); Glut, glutamate (n = 17); methyl, methylsulphonate (n = 10). Statistical inference One-way ANOVA with Holm-Sidak multiple comparisons tests. H) [Ca^2+^]_i_ when bath is sequentially washed, W1-5, using the same solution timings as employed in A and F. Statistical inference by One-way ANOVA (n = 15, f = 3).

To confirm that the decreases in [Ca^2+^]_i_ observed with Cl^−^ substitution were not artefacts through changes in bath height or flow rate that may arise on bath exchange, the effect of consecutive exchanges of the perifusate with normal control Hanks solution was tested. [Ca^2+^]_I_ did not decrease with this protocol (Fig. 4H; One-Way ANOVA).

The observation that substitution of bath Cl^−^ with methylsulphonate did not increase [Ca^2+^]_i_, is at odds with an activation of L-type VGCCs and subsequent increase in [Ca^2+^]_i_ we predicted to occur on depolarization of the adipocyte Vm with decreased [Cl^−^]_o_ (Bentley *et al.*, 2014). Similarly, the experiments in which gluconate was substituted for [Cl^−^]_o_ but where extracellular Ca^2+^ was maintained at 2.6 mM only prevented the decrease in [Ca^2+^]_i_ originally caused by anion chelation of this cation and did not show a depolarization-mediated increase in [Ca^2+^]_i_. To check that a constitutively Ca^2+^ influx pathway was actually present in visceral adipocytes from mice like that observed in rat (Fedorenko *et al.*, 2019*b*), the effects of extracellular Ca^2+^ addition, and verapamil a blocker of L-type VGCCs in these cells (Fedorenko *et al.*, 2019*b*) were tested.

Figure 5A shows that elevation of extracellular [Ca^2+^]_o_ to 5 mM reversibly increased [Ca^2+^]_i_ by 6% (2.3 to 9.8, 95% C.I.; Fig. 5B). In this particular experiment, subsequent substitution of extracellular Cl^−^ with gluconate to give 113, 53 and 18 mM [Cl^−^]_o_ (estimated [Ca^2+^]_o_ 1.24, 0.47 and 0.3 mM respectively with a K_a1_ of 17 L M^−1^) all decreased [Ca^2+^]_i_. However, in this particular experimental data set only 18 mM, the lowest [Cl^−^]_o_ tested, significantly lowered [Ca^2+^]_i_ relative to basal (Fig. 5B; Freidman, Dunn’s multiple comparison test). The magnitude of the decrease in [Ca^2+^]_i_ seen with gluconate (removal of bath Ca^2+^) was negatively correlated with the increase in [Ca^2+^]_i_ observed on elevation of bath Ca^2+^ (Spearman r = −0.33, p=0.05, n=35; Fig. 5C). It appears that the ability of an adipocyte to elevate [Ca^2+^]_i_ in response to an increased Ca^2+^ influx was mirrored by manipulations that decreased Ca^2+^ influx to diminish [Ca^2+^]_i_.

**Figure 5.**
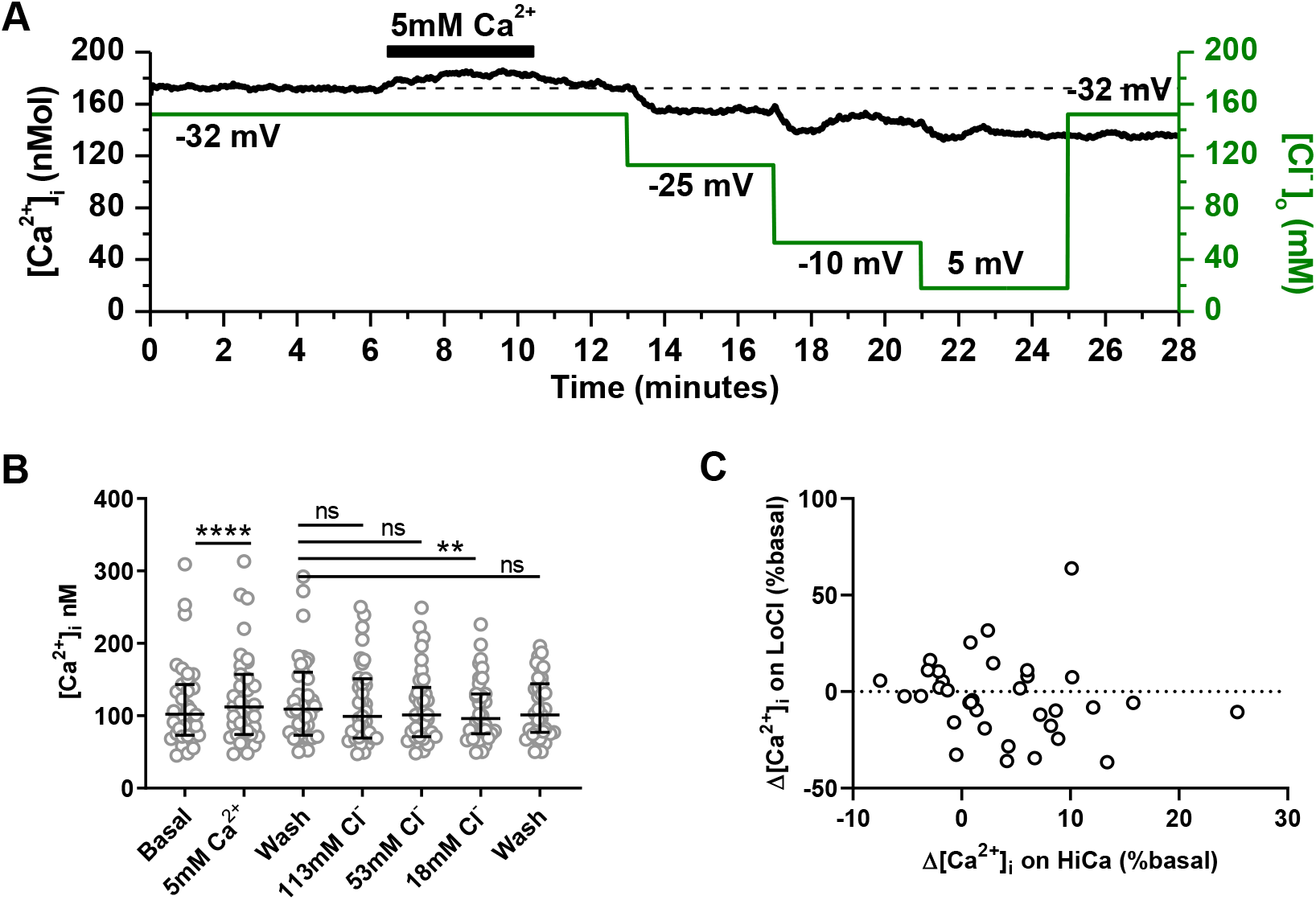
Changes in bath [Ca^2+^] are reflected in the intracellular [Ca^2+^] of white fat adipocytes. A) Mean intracellular Ca^2+^ for 6 adipocytes in response to an increase in bath [Ca^2+^], followed by sequential substitution of 39, 99 and 134 mM bath Cl^−^ with gluconate. Cells are all from the same field of view. B) [Ca^2+^]_i_ under the conditions indicated as shown in A) (n = 35, f = 9). Statistical inference by Friedman with Dunn’s multiple comparison tests. C) Relationship between the decrease in [Ca^2+^]_i_, Δ[Ca^2+^]_i_ on LoCa, observed with 134 mM gluconate and the increase in [Ca^2+^]_i_, Δ[Ca^2+^]_i_ on HiCa, observed on elevation of bath [Ca^2+^].

Figures 6A & B illustrate that analyses of all experiments in which extracellular Ca^2+^ was increased to elevate [Ca^2+^]_i_, [Ca^2+^]_i_ positively correlated with basal [Ca^2+^]_i_ (Spearman r = 0.22, p=0.04; Fig. 6A): cells with a higher basal apparently have greater sensitivity to increased Ca^2+^ influx. Conversely, the ability of a decrease in extracellular Ca^2+^ to diminish [Ca^2+^]_i_ (Fig. 6C) was negatively correlated with basal [Ca^2+^]_i_ (Spearman r = −0.26, p=0.04, n=63; Fig. 6D): cells with a higher basal apparently also have a greater sensitivity to a decreased Ca^2+^ influx. To check that the changes in [Ca^2+^]_i_ seen on elevation of [Ca^2+^]_o_ were not due to changes in surface charge mediated by the increase in divalent cation strength (Smith *et al.*, 1993), the effect of 2.5 mM MgCl_2_ addition to the bath solution was tested; this intervention was without effect (Wilcoxon, p =0.38, n=10). To confirm that the increase in [Ca^2+^]_i_ produced by an increase bath Ca^2+^ were mediated by L-type VGCCs like that observed in rat (Fedorenko *et al.*, 2019*b*), the effect of verapamil was tested. Figure 6E shows that 20 μM verapamil, relative to that seen with the 0.1% DMSO vehicle control, abolished the ability increased [Ca^2+^]_o_ to elevate [Ca^2+^]_i_ (p<0.001, Kruskal Wallis, Dunn’s multiple comparison test). Figure 6F shows that at 20 μM verapamil decreased basal [Ca^2+^]_i_ relative to that observed with the 0.1% DMSO vehicle control (p=0.029, Kruskal Wallis, Dunn’s multiple comparison test). However 10 μM BAY-K8644, an L-type VGCC agonist, failed to affect [Ca^2+^]_i_ (p=0.3, Kruskal Wallis, Dunn’s multiple comparison test).

**Figure 6.**
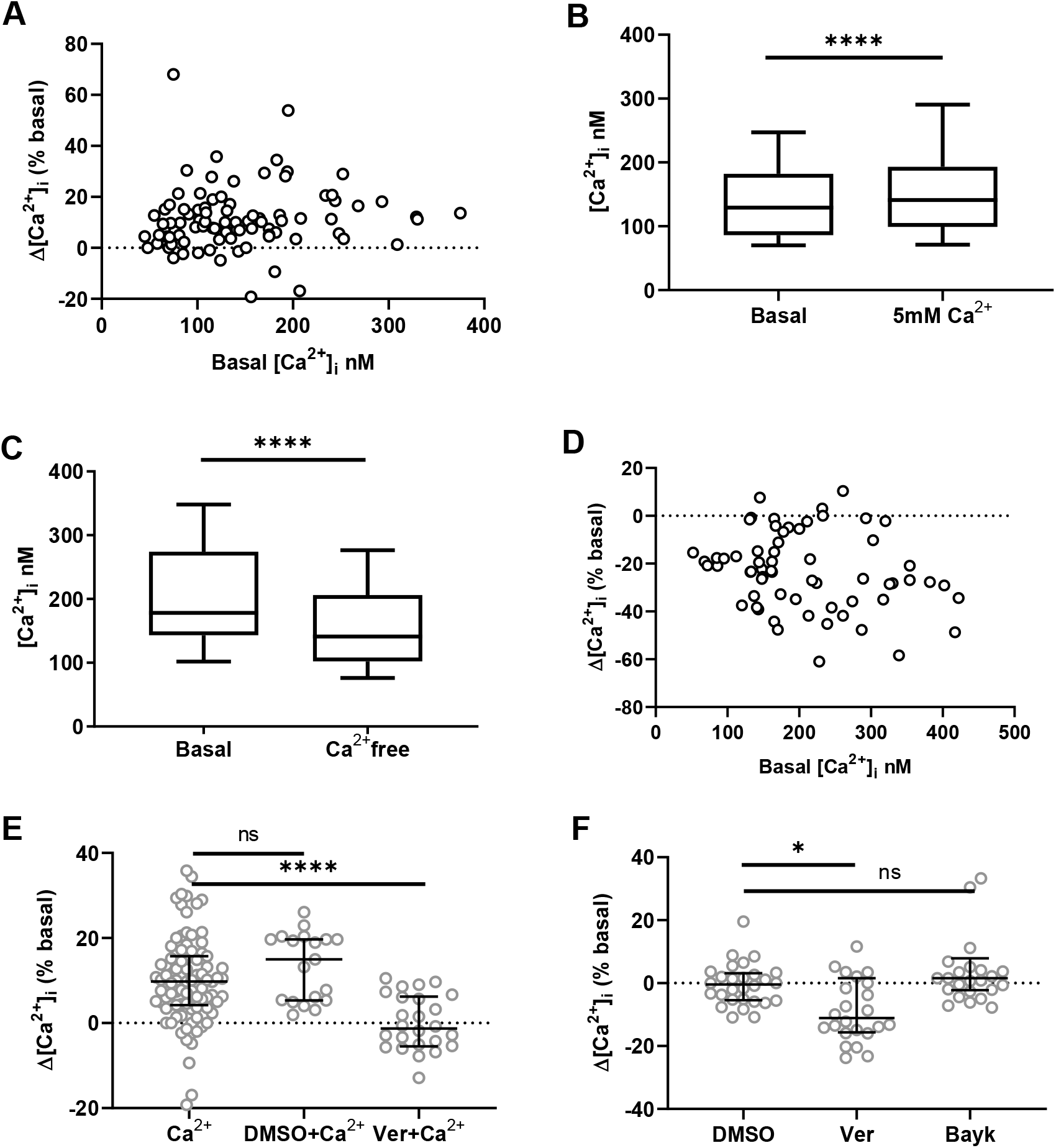
Verapamil inhibits Ca^2+^ influx. A) Relationship between the increase in [Ca^2+^]_i_, Δ[Ca^2+^]_i_, observed on elevation of bath [Ca^2+^] with basal [Ca^2+^]_i_. B) Effect of an increase in [Ca^2+^]_o_, 5 mM Ca^2+^, compared to basal [Ca^2+^]_i_ (n = 91, f = 23). Statistical inference by Wilcoxon. C) Effect of a decrease in [Ca^2+^]_o_, Ca^2+^ free, compared to basal [Ca^2+^]_i_ (n = 63, f = 9). Statistical inference by Wilcoxon. D) Relationship between decrease in recovery in [Ca^2+^] Δ[Ca^2+^]_i_, observed on removal of bath [Ca^2+^] with basal [Ca^2+^]_i_. E) Change in [Ca^2+^]_i_, Δ[Ca^2+^]_i_, observed in response to an elevation of bath Ca^2+^ with: no other addition, Ca^2+^ (n = 91); in the presence of 0.1% DMSO, DMSO+Ca^2+^ (n = 19); and in the presence of 0.1% DMSO + 20 μM verapamil (n = 26). Statistical inference by Kruskal-Wallis with Dunn’s multiple comparison tests. F) Change in [Ca^2+^]_i_, Δ[Ca^2+^]_i_, in 2.5 mM [Ca^2+^]_o_ observed in response to: 0.1% DMSO, DMSO (n = 28); 0.1% DMSO + 20 μM verapamil, Ver (n = 24); and 0.1% DMSO + 10 μM BAY-K8644 (n = 26). Statistical inference by Kruskal-Wallis with Dunn’s multiple comparison tests.

As electrochemical depolarization of Vm did not promote L-type VGCC activity and an increase in [Ca^2+^]_i_, the effect of growth hormone was explored since this has been deomsontrated to elevate [Ca^2+^]_i_ in white fat adipocytes from rat (Gaur *et al.*, 1996, 1998).

Figure 7A shows that after a delay of several minutes, 10 nM recombinant human growth hormone, GH, elevated [Ca^2+^]_i_. This change in [Ca^2+^]_i_ was manifest either as a transient increase, a maintained increase, or as oscillations. Out of 11 cells tested 64% responded to 10 nM GH (f = 3). Similar results were also seen with 20 nM GH, where out of 54 cells tested, 48% responded with an elevation in [Ca^2+^]_i_ (Fig. 7B). Responses to GH were abolished on removal of extracellular Ca^2+^ (n = 70, f = 11, a = 5; Fig. 7D); data which suggest the increase in [Ca^2+^]_i_ seen with GH is due to Ca^2+^ influx and not mobilization from internal stores. Furthermore, 20 μM verapamil also abolished GH evoked increases in [Ca^2+^]_i_ (n =22, f = 5, a = 3; Figs. 7C, D); data which supports the idea that GH promotes L-type VGCC activity to elevate [Ca^2+^]_i_ (Gaur *et al.*, 1996, 1998). In 53 cells tested, addition of 0.4% vol/vol water, the vehicle for GH, did not elicit Ca^2+^ responses (f = 15, a = 7s; Fig. 7D).

**Figure 7.**
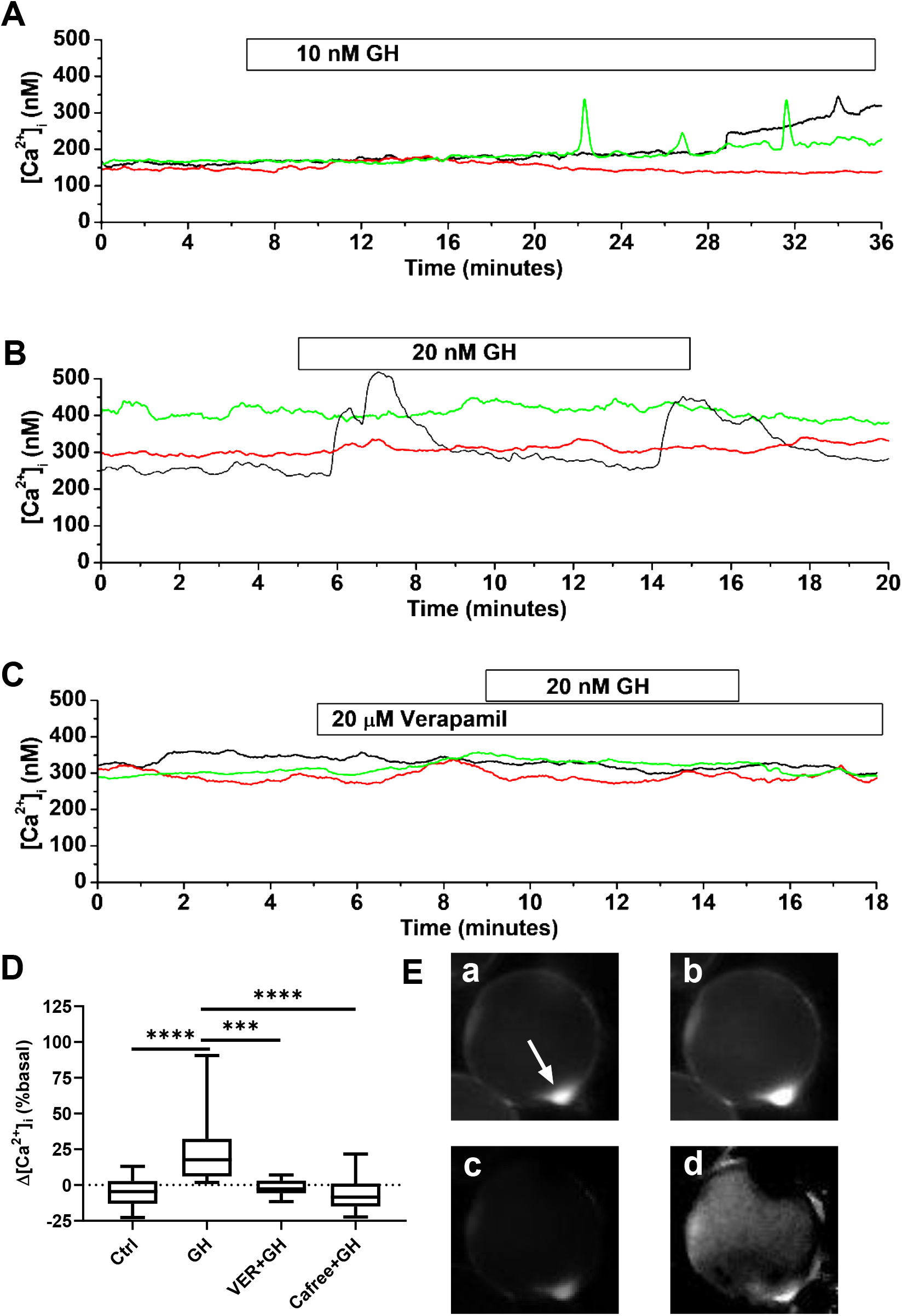
Growth hormone stimulates Ca^2+^ influx. Representative traces of intracellular Ca^2+^, [Ca^2+^]_i_, for white fat adipocytes under the conditions indicated. A) [Ca^2+^]_i_ in 3 adipocytes in response to addition of 10 nM growth hormone (GH). Cells are all from same field of view. B) [Ca^2+^]_i_ in 3 adipocytes in response to addition of 20 nM growth hormone (GH). Cells are all from same field of view. C) [Ca^2+^]_i_ in 3 adipocytes in response to addition of 20 nM growth hormone (GH) but in the continued presence of 20 μM verapamil. Cells are all from same field of view. D) Change in [Ca^2+^]_i_, Δ[Ca^2+^]_i_, observed in response to: the peptide vehicle control 0.2% H_2_O_2_, Ctrl (n = 30); 10 or 20 nM Growth Hormone, GH (n = 60); 20 nM Growth hormone in either the presence of 20 μM verapamil, VER+GH (n=15) or absence of extracellular Ca^2+^, Cafree+GH (n = 61). Both responders and non-responders are included for all treatments. Statistical inference by Kruskal-Wallis with Dunn’s multiple comparison tests between all treatments, non-significant differences are not shown. E) Grayscale epifluorescent Ca^2+^ images of a field of a single mouse epididymal white fat adipocyte taken from A) above. a) under basal, b) in response to 20 nM growth hormone and cb), the difference between images a) and b). Image are 75 μm square and are the averages of 50 frames; images a, b and c have identical brightness and contrast. Note nuclear protuberance, arrowed, and the brighter circumferential fluorescence where the cytoplasm has the deepest volume parallel to the plane of illumination. d) Ratio of image ii and i, to normalize dye loading and cytoplasmic volume. Note the particularly bright fluorescence in the perinuclear region.

Figure 7E demonstrates that the increase in [Ca^2+^]_i_ evoked by 20 nM GH appears to originate in the perinuclear region of the adipoctye; a similar observation was made for all adipocytes who responded to GH and for which the nuclear knob could be identified (n > 5).

## Discussion

We have shown that a variety of membrane impermeant organic anions that are commonly used to substitute for Cl^−^, chelate free Ca^2+^ to directly effect [Ca^2+^]_i_ of the cells under study. Compared to gluconate, aspartate and glutamate, methylsulphonate had the lowest association with Ca^2+^ and was the best substitute for bath Cl^−^. A similar conclusion was made previously for patch-pipette solutions (Woehler *et al.*, 2014). The rank order for association of gluconate > aspartate > glutamate > methylsulphonate we report is comparable to that deduced from published values (Table 2). In previous studies, the association constant for metal cation chelation was either derived from the change in its acid dissociation constant,pKa, (Lumb & Martell, 1970) or measured directly by Ca^2+^ electrodes (Vavrusova & Skibsted, 2013; Vavrusova *et al.*, 2013). Although we deduced the stoichiometry of Ca^2+^ chelation by statistical methods, all other reports have assumed a 1:1 for simplicity (Table 2). This has implications for gluconate for where a binary 2:1 complex was favoured over 1:1. With a 1:1 model and using equation 8 we reevaluate the K_a_ for gluconate as 37±7.4 L M^−1^ (n=4) (Table 2).

**Table 2.** Primary association constant, K_a_ (L M^−1^) for Ca^2+^ anion salts with a 1:1 stoichiometry. A, present study (K_a_ for gluconate in parentheses re-evaluated for a 1:1 binding); B, I = 0.1 25°C via pKa (Lumb & Martell, 1970); C, I = 0.2 25°C via ECa (Vavrusova & Skibsted, 2013; Vavrusova *et al.*, 2013); D, I = 0.1 25°C via pKa (Skibsted & Kilde, 1972); E, I = 0.2, 25°C via pKa (Cannan & Kibrick, 1938); F, I=0.2 25°C via pKa (Schubert & Lindenbaum, 1952). G, I = 0.18 21-24°C via ECa, K_a_ values derived by fits of equation to [Ca derived from quoted ratios shown in Table 1 of Kenyon and Gibbons (Kenyon & Gibbons, 1977). Note methylsulphonate is a synonym for methanesulphonate.

| Anion | A | B | C | D | E | F | G |
| --- | --- | --- | --- | --- | --- | --- | --- |
| Gluconate | 17 (37) |  | 31 | 32 | 16 | 17 | 33 |
| Aspartate | 12 | 40 | 7 |  |  |  |  |
| Glutamate | 8.2 | 27 | 3 |  |  |  |  |
| Methylsulphonate | 3.3 |  |  |  |  |  | 0.8 |

Our K_a_ values are comparable to those previously determined with 1:1 binding coordination and equivalent ionic strengths. It should be noted from table 2 that K_a_ is larger when I is small (Lumb & Martell, 1970) an observation consistent with Debye-Hückel theory. The range of published K_a_ values for any given anion varies up to 2-fold and most likely relates to differences in experimental conditions and assumptions made in the separate studies. The source of variation behind our K_a_ values was not obvious given the controls undertaken, but should be noted when estimates of free Ca^2+^ are calculated using the K_a_ values presented here.

The ability of membrane impermeant glutamate and gluconate to chelate bath Ca^2+^ readily explains why substitution of bath Cl^−^ with these anionic species decreased adipocyte [Ca^2+^]_i_. This action, mimicked by the substitution of bath Ca^2+^ with Mg^2+^ and abrogated by the addition of extra Ca^2+^ to the bath solution to replace that sequestered by the anion, support this idea. Although not the first to note the adverse effects of organic anions on biological phenomena due to Ca^2+^ chelation (Christoffersen & Skibsted, 1975; Kenyon & Gibbons, 1977) we do now provide K_a_ values determined in solutions of physiological ionic strength and pH. These values allow estimation of [Ca^2+^]_i_ to help explain anomalous observations when these organic anions are used in physiological experiments. For example, since glutamate chelates Ca^2+^ we can estimate the free concentration of the anion when it is applied as an ionotropic neurotransmitter. Interestingly, the decrease in [Ca^2+^]_o_ and [glutamate^−^]_o_ due to chelation is minimal: if we take K_a_ as 0.0082 L mM^−1^ (table 2) then [Ca^2+^]_o_ is estimated to drop from 2.5 to 2.498 mM and [glutamate^−^]_o_ from 100 to 98 μM. However, the chelation of Ca^2+^ by anions becomes more of a problem when Cl^−^ is substituted by organic anions on the magnitude of 10-100 mM as we show here with extracellular solutions and by others with intracellular patch pipette solutions (Woehler *et al.*, 2014).

Although the Vm of murine visceral white fat adipocytes is dependent on their plasma membrane permeability to Cl^−^ (Bentley *et al.*, 2014), and they possess a basal Ca^2+^ influx mediated by constitutive activity of L-type VGCCs (Fedorenko *et al.*, 2019*b*), the manipulation of extracellular Cl^−^ by substitution with methylsulphonate failed to affect [Ca^2+^]_i_. This conundrum was not due to the absence of L-type VGCCs activity since verapamil, a potent blocker of these ion channel species, could decrease both basal and potentiated [Ca^2+^]_i_ in accord with this model (Fedorenko *et al.*, 2019*b*). Moreover, this model is supported by our observation that mouse white fat adipocytes express the transcripts of both CaV1.2 and CaV1.3 voltage-gated calcium channels (Fedorenko *et al.*, 2019*a*) and have a ^45^Ca^2+^ influx sensitive to L-type VGCC antagonists (Martin *et al.*, 1975). Unfortunately, BAY-K 8644 a dihydropyridine agonist of this channel class did not readily promote basal Ca^2+^ influx and elevate [Ca^2+^]_i_. The reasons for this are unclear, but may relate to interference with the dihydropyridine binding to L-type VGCC by free fatty acids (Pepe *et al.*, 2006) and argued elsewhere (Fedorenko *et al.*, 2019*b*).

Previous results produced in white fat adipocytes with substitution of [Cl^−^]_o_ by aspartate (Bentley *et al.*, 2014) should be interpreted with caution, not just because of extracellular Ca^2+^ chelation but also due to partial permeability (Pulbutr *et al.*, 2007).

Since, the magnitudes of the elevation and depression in [Ca^2+^]_i_ observed with increased and decreased extracellular Ca^2+^ respectively were correlated with the level of basal [Ca^2+^]_i_ further supports the idea that basal [Ca^2+^]_i_ reflects the ratio of Ca^2+^ influx and efflux. Where removal of bath Ca^2+^ by chelation or Mg^2+^ substitution had a larger effect in adipocytes with higher basal [Ca^2+^]_i_, data that proposes that these particular cells either have a greater prevalence of L-type VGCC activity or/and lower activity of Ca^2+^ extrusion than adipocytes with lower basal [Ca^2+^]_i_.

Why L-type VGCCs failed to respond to Cl^−^ mediated changes in Vm may relate to novel gating characteristics of these channels. It appears hormonal, and not Vm, is the predominant factor that regulates activity of this channel moiety in adipocytes; for example GH as shown here and by others (Gaur *et al.*, 1996, 1998). Teleologically, slow biochemical modulation of ion channels is apt for a cell type not required to respond on a sub-second timescale, unlike electrically excitable cells. Indeed, changes in expression level are seen in white fat adipocytes with aging (Fedorenko *et al.*, 2019*a*) and in response to parathyroid hormone (Ni *et al.*, 1994) and growth hormone (Gaur *et al.*, 1998). Why L-type VGCCs and their associated Ca^2+^ influx is concentrated to the perinuclear region of white fat adipocytes, and what the physiological consequences of this and its regulation are, have yet to be explored.

### Declaration of interest, funding and acknowledgements

The authors declare that there is no conflict of interest that could be perceived as prejudicing the impartiality of the research reported. NA is in receipt of a Schlumberger Foundation PhD fellowship. PAS wishes to thank the Leverhulme Trust (Grant ID: RPG-2017-162) for financial support. NA conducted and analyzed the experiments, PAS designed the experiments, performed the modelling and statistics, and wrote the manuscript.

Ca2+ chelation by organic anions reveal novel calcium channel gating

